# A one-shot learning signal in monkey prefrontal cortex

**DOI:** 10.1101/2020.11.27.401422

**Authors:** Jascha Achterberg, Mikiko Kadohisa, Kei Watanabe, Makoto Kusunoki, Mark J Buckley, John Duncan

**Author notes:** Correspondence: John Duncan, MRC Cognition and Brain Sciences Unit, 15 Chaucer Road, Cambridge CB2 7EF, UK.

## Abstract

Much animal learning is slow, with cumulative changes in behavior driven by reward prediction errors. When the abstract structure of a problem is known, however, both animals and formal learning models can rapidly attach new items to their roles within this structure, sometimes in a single trial. Frontal cortex is likely to play a key role in this process. To examine information seeking and use in a known problem structure, we trained monkeys in a novel explore/exploit task, requiring the animal first to test objects for their association with reward, then, once rewarded objects were found, to re-select them on further trials for further rewards. Many cells in the frontal cortex showed an explore/exploit preference, changing activity in a signal trial to align with one-shot learning in the monkeys’ behaviour. In contrast to this binary switch, these cells showed little evidence of continuous changes linked to expectancy or prediction error. Explore/exploit preferences were independent for two stages of the trial, object selection and receipt of feedback. Within an established task structure, frontal activity may control the separate operations of explore and exploit, switching in one trial between the two.

**Significance statement:** Much animal learning is slow, with cumulative changes in behavior driven by reward prediction errors. When the abstract structure a problem is known, however, both animals and formal learning models can rapidly attach new items to their roles within this structure. To address transitions in neural activity during one-shot learning, we trained monkeys in an explore/exploit task using familiar objects and a highly familiar task structure. In contrast to continuous changes reflecting expectancy or prediction error, frontal neurons showed a binary, one-shot switch between explore and exploit. Within an established task structure, frontal activity may control the separate operations of exploring alternative objects to establish their current role, then exploiting this knowledge for further reward.

---

Much animal learning occurs slowly, with prediction errors leading to incremental changes in the link between actions and their outcomes (1, 2). A similar process of incremental change in response to experience underlies powerful learning models, from early versions of backpropagation to modern deep learning systems (3, 4). Animals and formal models are also capable, however, of rapid, sometimes one-shot learning. When the abstract structure or schema of a problem is known, new items can rapidly be attached to their roles within this structure (5). Familiar examples include learning to learn (6), object-location binding (7) and meta-learning (8). One-shot binding of items to roles is conspicuous throughout human cognition, endowing thought and behavior with their characteristic speed, flexibility and compositionality (9).

Frontal cortex is thought to play a central role in binding together the components of a cognitive operation (10, 11), likely as part of a broader cognitive control network (12, 13). In the behaving monkey, many aspects of a current task are represented in the firing of frontal neurons (14), with prominent “mixed selectivity” or conjunctive coding (15, 16) of different task features. Conjunctive coding is a likely substrate for variable binding, allowing the components of a cognitive operation to be correctly combined (10, 17). Frontal neurons are well known to encode trial-specific instructions and working memory items, indicating how individual decisions should be taken within a well-learned task structure (18–20). In a block of trials, frontal population activity shows abrupt changes when new task rules are adopted (21) or object-reward bindings must be reversed (22).

In an important series of studies (23–26), Procyk and colleagues used a spatial selection task to examine rapid transition from unknown to known rules. In this task, monkeys selected different screen locations in turn, searching for the one location associated with reward (“explore” trials). Once reward was found, the same location could be selected on a series of further trials (“exploit” trials) for further rewards. Monkeys performed this task close to perfectly, with immediate transition from explore to exploit once the rewarded location was found. At this transition, spatial selectivity declined in neurons of dorsolateral frontal cortex ((23); though see (26)), and response to feedback decreased in anterior cingulate neurons (24). To extend this investigation of rapid learning, we used a similar comparison of explore and exploit trials in an object selection task. Reflecting the different computations required at object selection and receipt of feedback, we examined explore/exploit preferences at these different task phases. We focus primarily on lateral prefrontal cortex, with comparison data from inferior parietal cortex.

On each trial (Figure 1A), monkeys selected (by touching) one object from a set of four presented on a touchscreen, bringing reward or nonreward. In each session, the animal worked through a series of problems, based on two fixed sets of 4 objects (Figure 1A) which alternated between problems. The choice display for each trial contained all four objects of the current set, randomly positioned. For each new problem, either one or two objects from the current set were targets, bringing reward when touched, while the remainder were non-targets bringing no reward. Each problem, accordingly, required new object-role bindings to be learned. On a first cycle of trials (explore), the monkey sampled objects in turn across a series of trials, searching for the rewarded target or targets. 1-target and 2-target problems were blocked, so the animal knew in advance how many to discover. Once targets were found, there followed 3 exploit cycles, in which animals were rewarded for re-selecting the targets discovered in cycle 1. In 2-target problems, the animal was free to select the two targets in each cycle in either order. Each target was rewarded just once per cycle, and all cycles ended as soon as the single target or two targets had been selected. Optimally, therefore, the explore cycle consisted of a random sequence of object selections, avoiding re-visits, until the single target (mean expected number of trials = 2.50) or two targets (mean expected number of trials = 3.33) were discovered. Exploit cycles consisted optimally of just one (1-target problems) or two (2-target problems) trials. We reasoned that explore trials called for a process of information seeking guided by long-term knowledge of task structure – selecting candidate objects from the available set, and based on reward or nonreward, linking these objects to their roles as target or non-target. Exploit trials, in contrast, called for use of these newly-learned roles, selecting targets and avoiding non-targets. We recorded neural activity in lateral prefrontal cortex, within and surrounding the principal sulcus (Figure 1B), searching for sharp changes in activity between explore and exploit phases.

**Figure 1.**
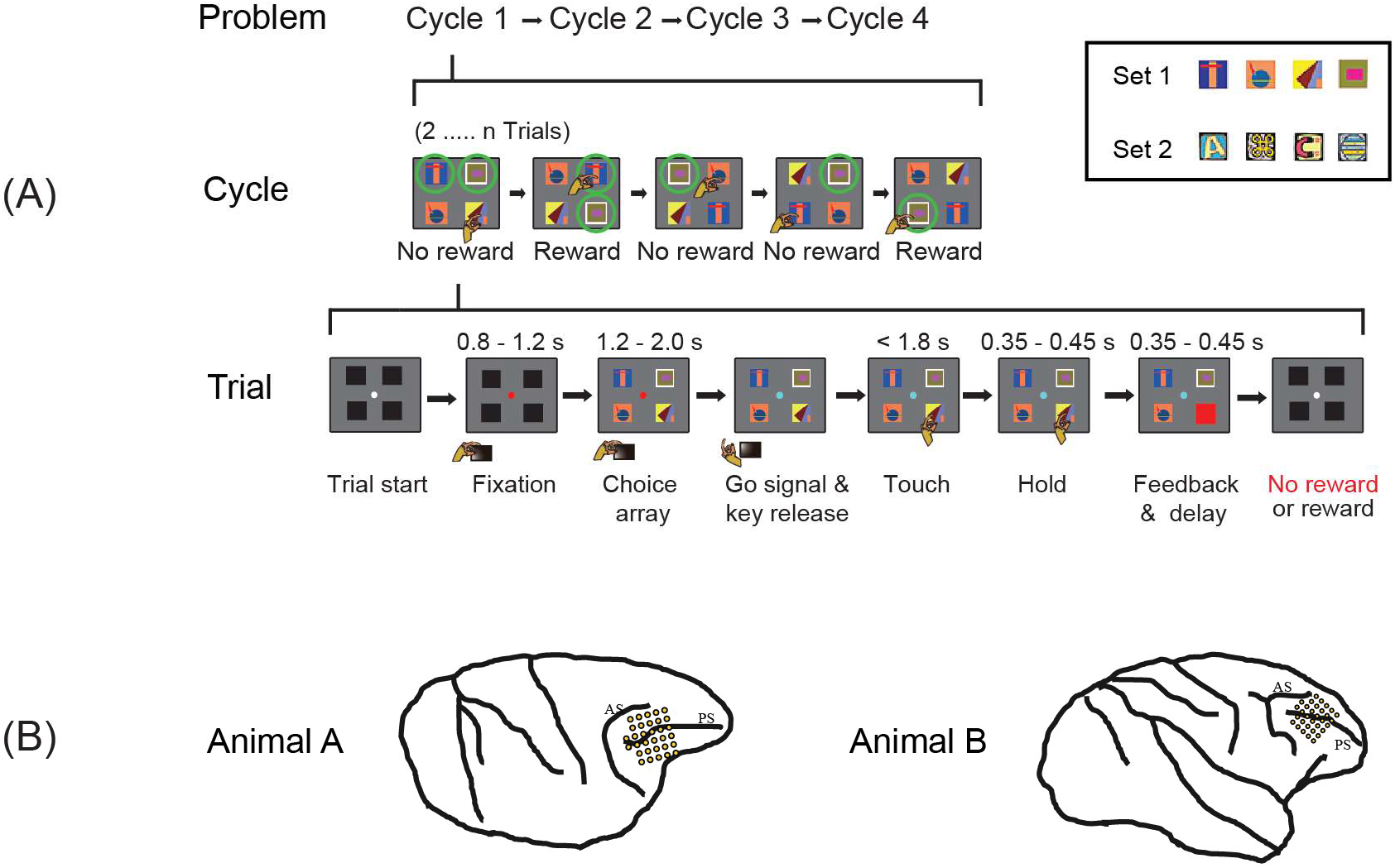
(A) Object selection task, 2-target version. On each trial, the monkey touched a single object in a visual display. For each new problem, in the first set of trials (cycle 1), the monkey selected one object after another, learning which 2 objects (targets) were associated with reward. Targets are indicated here by green circles (not present on actual display). In subsequent trials (cycles 2-4), the animal could reselect the same targets for further rewards. After 4 cycles, targets were redefined for the next problem. Alternate problems used two different 4-object sets, fixed for each animal (object sets for one animal in inset). Equivalent 1-target problems had only a single target. (B) Recording areas in each animal. PS – principal sulcus; AS – arcuate sulcus.

Human brain imaging suggests that first encounters with a new problem lead to strong activity in lateral frontal cortex and other cognitive control regions, which rapidly decreases once the solution is found (27, 28). At the level of single neurons, we report bidirectional changes, with some cells selectively active during explore, shifting in one trial to others selectively active during exploit.

## Results

### Behavior

Behavioral data are summarized in Figure 2, separately for 1-target (left column) and 2-target (right column) problems. Data are averages for the two animals; individual data are shown in SI Appendix, Figures S1-2. In 1-target problems, the mean number of trials per cycle was close to optimal (Figure 2A, left; data in red, optimal possible performance in blue), indicating rapid, generally one-trial learning. In 2-target problems (Figure 2A, right), performance improved more gradually over cycles. A more detailed breakdown of response types is shown in Figure 2B. In each cycle, the number of correct target selections (red) was by definition one (1-target problems) or two (2-target problems). As expected, novel non-target selections (selection of a non-target not previously sampled in this cycle) were frequent in cycle 1, occurring in the proportions required by a random search. Rapid discrimination between targets and non-targets is shown by the substantial decline in non-target selections between cycles 1 and 2, clearly evident in both 1-target and 2-target problems. Revisits to an object already sampled in a cycle were infrequent throughout (non-target revisits – acqua; target revisits – purple, impossible for 1-target problems). A further analysis of cycle 1 data (Table 1) confirms this strong avoidance of objects already sampled. For this analysis, cycle 1 trials were broken down according to the number of objects already sampled, from 0 (first trial of cycle) to 3. The table shows mean percentage of trials with revisit to a previously sampled object, compared to expected values for a random selection (see SI Appendix, Table S1 for individual data). In all cases, revisit percentage was far below the chance expectation. Together, these data confirm high-speed learning in the task, with strong avoidance of objects already sampled within a cycle, and from cycle 2 onwards, excellent discrimination between targets and non-targets, especially in 1-target problems.

**Table 1.** Cycle 1: Percentage of trials with re-selection of a previously sampled object, as a function of number already sampled.

| Already sampled | Data |  | Expected |
| --- | --- | --- | --- |
|  | *1-target problems* | *2-target problems* |  |
| **0** | 0% | 0% | 0% |
| **1** | 4.6% | 5.9% | 25% |
| **2** | 19.1% | 20.7% | 50% |
| **3** | 48.1% | 49.1% | 75% |

**Figure 2.**
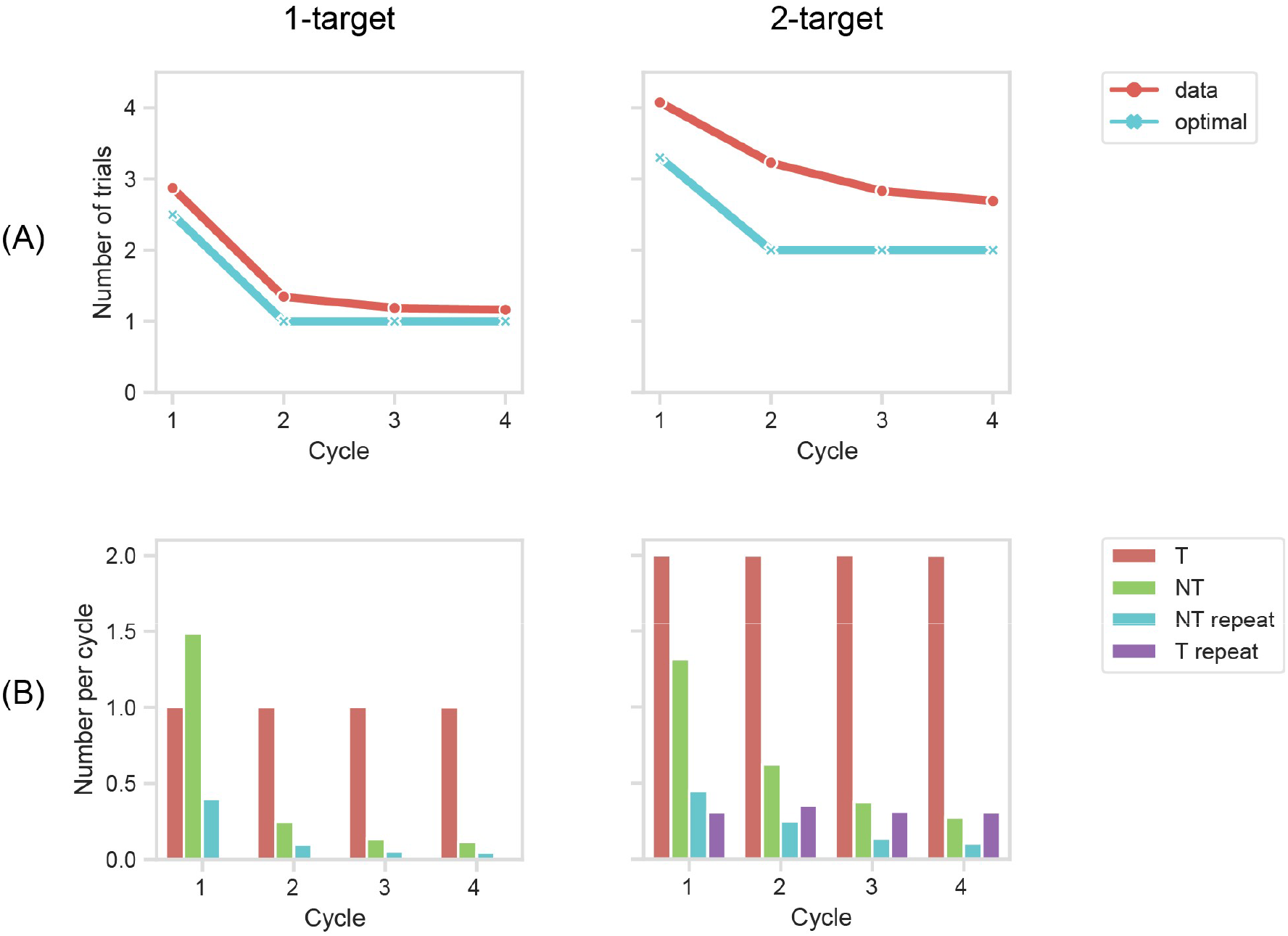
Behavioral data. (A) Mean number of trials per cycle. (B) Trials per cycle broken down into correct target selections (T), novel non-target selections (selection of a non-target not previously sampled in this cycle, NT), repeat non-target selections (selection of a non-target previously sampled in this cycle, NT repeat), and repeat target selections (T repeat; only possible for 2-target problems).

### Prefrontal cells show preference for explore or exploit

Across 56 task sessions, we recorded activity from 254 cells, 176 from monkey A and 78 from monkey B, in a region spanning the principal sulcus and adjacent dorsolateral and ventrolateral frontal convexities. Except where otherwise specified, data were analyzed just from correct trials, i.e. those on which a current target object was selected.

For our first analysis we asked whether frontal neurons differentiate the processes of explore – seeking now information to bind into the problem structure – vs exploit – using known information to guide behavior. To give the strongest measure of explore/exploit preferences, we focused initially on a comparison of cycles 1 and 4, combining data from the rapidly-learned 1-target problems and the more slowly learned 2-target problems. As the structure of the task requires quite different computations at different stages of the trial, we analyzed data from two trial phases: choice (CH), the period following onset of the choice display, and feedback (FB), the period following onset of feedback. To ensure unbiased results, we adopted a cross-validated approach. For each cell, trials were randomly split into two halves. On the first half of the data, we performed ANOVA with factors cycle (1, 4) x number of targets (1, 2) x object set x touched location (1-4). These ANOVAs used data from two 400 ms windows, beginning at onset of CH and FB, with a separate ANOVA for each window. For each analysis window, cells with a significant (P < .05) main effect of cycle were classified as “explore” (spike rate cycle 1 > 4) or “exploit” (spike rate cycle 1 < 4). For each explore or exploit cell, we extracted peri-stimulus time histograms (PSTHs) from the other half of the data. For a more complete view of the data, these unbiased PSTHs extended across a longer period (−200 to +500 ms from event onset). PSTHs for each cell were normalized and then averaged across cells within each group (see Materials and Methods). T-tests across cells, again using 0-400 ms windows, were used to confirm significant cycle preference in the held out data.

For the CH period, the main effect of cycle was significant in 44 cells (17.3% of total), 18 with a preference for cycle 1, 26 for cycle 4. Cross validated mean PSTHs for these cells are shown in the upper row of Figure 3, with explore (cycle 1 preferring) cells in the left panel, exploit (cycle 4 preferring) cells on the right. T-tests on the window 0-400 ms from CH onset confirmed significant cycle preferences in these held out data (for explore cells, T = 4.58, P < .001; for exploit cells, T = 4.39, P < .001). PSTHs suggest sustained cycle preferences that began even before CH onset. Notably, however, cells selected for a cycle preference during CH showed no evidence of a similar preference around FB.

**Figure 3.**
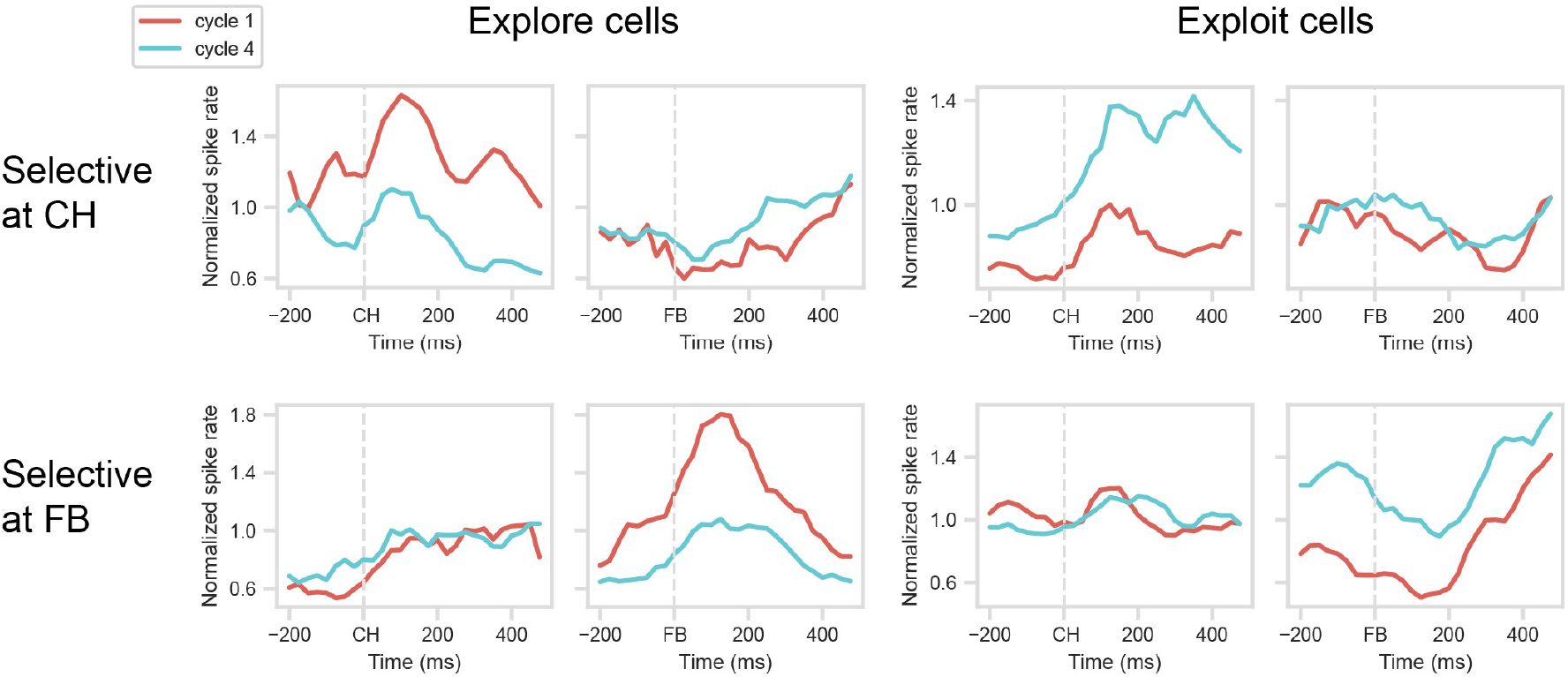
Mean normalized spike rate for explore (spike rate cycle 1 > cycle 4) and exploit (spike rate cycle 4 > cycle 1) cells, identified at CH and FB periods. Note that data are cross-validated, with separate trials used to identify selective cells and to construct PSTHs.

A similar picture is evident for cells with cycle preference at FB (Figure 3, lower row). ANOVA on the FB period showed a significant main effect of cycle in 52 cells (20.5% of total), 15 with a preference for cycle 1, 37 for cycle 4. PSTHs suggest sustained cycle preferences around FB, with T-tests on the held out data confirming significant cycle preferences in the 0-400 ms window (for explore cells, T = 2.94, P < 0.05; for exploit cells, T = 6.61, P < 0.001). Again, however, cells selected for a cycle preference during FB showed no evidence of a similar preference around CH.

These initial analyses show that substantial fractions of prefrontal cells differentiate the processes of explore and exploit. Though explore/exploit preferences are seen at both CH and FB, preferences at these two stages of a trial are unrelated, implying selectivity for the conjunction of cycle (explore/exploit) and trial stage (CH/FB).

### Temporal cross-generalization of cycle preferences

To confirm that cycle preference is stable within a task phase (CH, FB) but not across task phases we used a temporal cross-generalization analysis. We again split the data for each cell into two halves, and for each half, subtracted mean activity in cycle 1 from mean activity in cycle 4. We de-weighted the means to be unbiased for touched location (1-4) and number of targets (1, 2). For this analysis we used 100 ms windows, four windows from onset of CH and four from onset of FB. For each window, this produced 2 vectors of 254 cycle 4-1 differences, one for each half of the data, where 254 is the number of recorded cells. Correlations between vectors from the two halves of the data are shown in Figure 4. Strong correlations within CH and FB periods show that, within each period, the preference for cycle 1 vs 4 was stable; between periods, however, correlations close to zero show unrelated cycle preferences.

**Figure 4.**
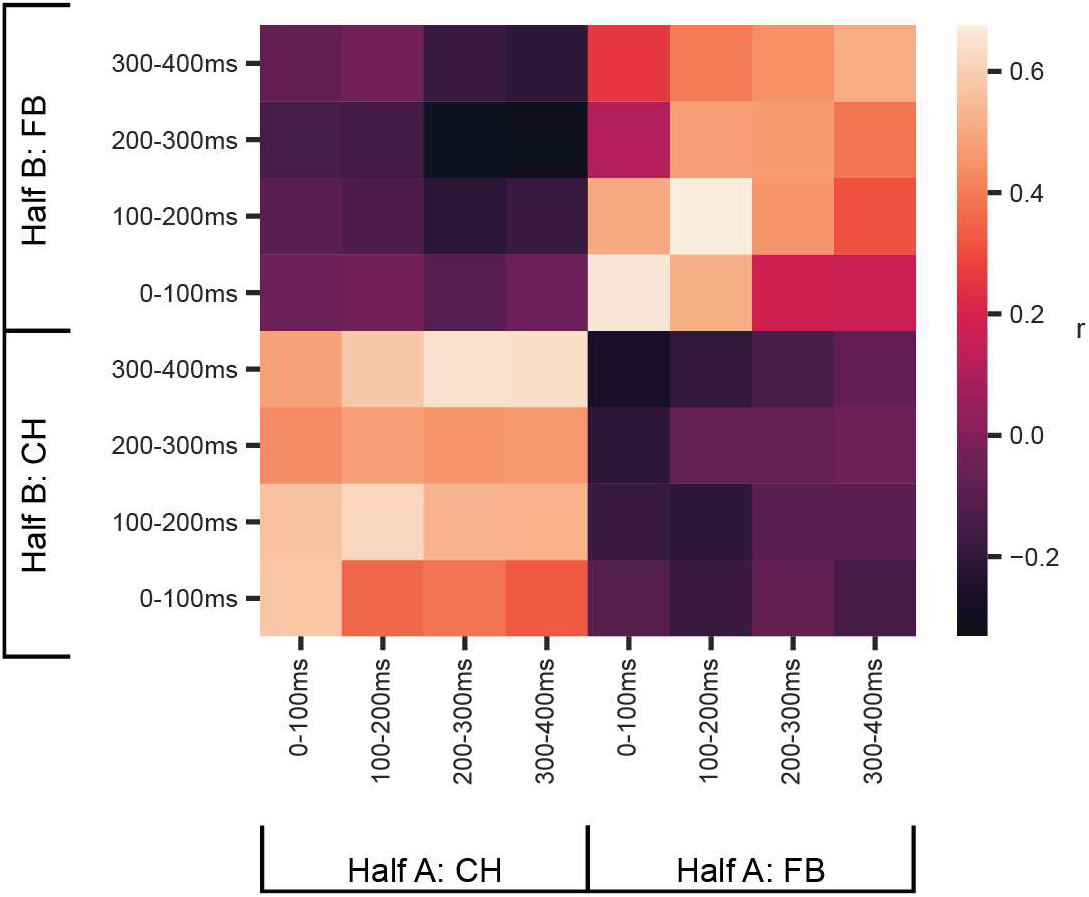
Temporal stability of explore-exploit preference. For each cell, trials were split into two halves (A and B), and within each half, cycle preference (cycle 4 spike rate minus cycle 1 spike rate) was calculated in 4 x 100 ms windows following onset of CH and FB. For each time window, this produced two independent vectors of 254 cycle preferences (1/cell). Data are correlations between half A and half B vectors.

### One-shot learning

Having established that patterns of frontal activity differentiate explore and exploit, we moved on to examine the transition between these patterns with rapid learning. For this purpose we focused on the one-shot learning seen in most 1-target problems. To mirror this rapid change in behavior, we searched for a similar one-shot change in the activity of explore/exploit cells. We focused on the explore and exploit cells identified in our first analysis, and examined their detailed behavior during the rapid learning of 1-target problems.

For the main analysis we compared activity across cycles 1 to 4. To focus on successful rapid learning, for cycles 2 to 4 we excluded the exceptional cases in which the first response of the cycle was incorrect. Mean PSTHs for the same 4 groups of cells were calculated as before, using just half the data for cycles 1 and 4 (the half not used in cell selection), but all data for cycles 2 and 3. Results are shown in the solid lines in Figure 5. Across the 4 cell groups, the results were clearcut, with activity on cycle 2 immediately switching from the cycle 1 to the cycle 4 pattern. For cells with an explore (cycle 1) or exploit (cycle 4) preference at CH (Figure 5, top row), tests on the 400 ms window following CH onset showed significant differences between cycle 1 and 2 (explore cells, T = 3.91, P = 0.001; exploit cells, T = 4.17, P = 0.001), but no significant differences between cycles 2 to 4 (explore cells, F = 0.03; exploit cells, F = 1.08). For cells with an explore (cycle 1) or exploit (cycle 4) preference at FB (Figure 5, bottom row), similar results were obtained in the 400 ms window following FB onset. The difference between cycles 1 and 2 approached significance for explore cells, T = 2.06, P = 0.058, and was significant for exploit cells, T= 7.54, P < 0.001, while differences between cycles 2 to 4 were not significant (explore cells, F = 2.26; exploit cells, F = 0.25).

**Figure 5.**
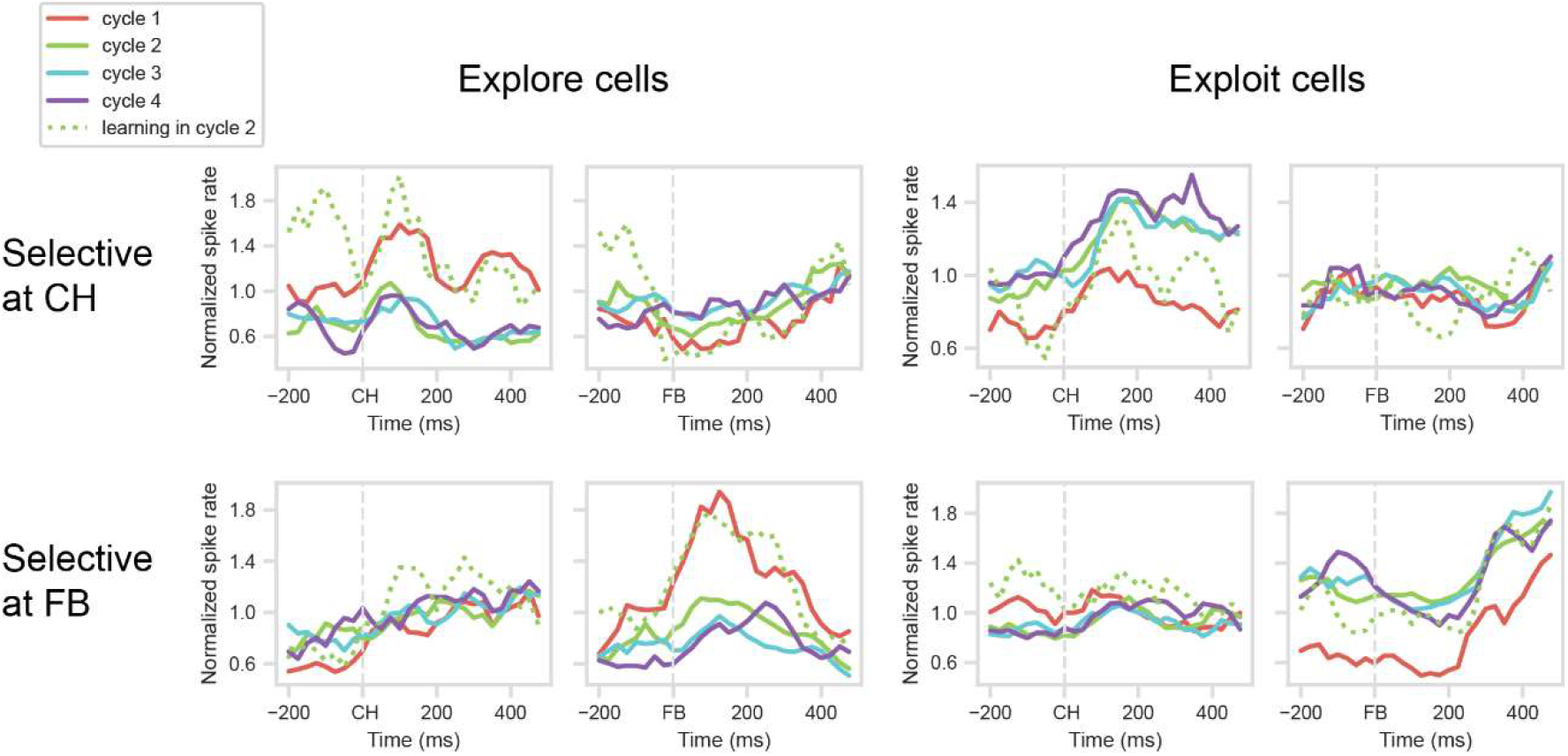
1-target-problems: Mean normalized spike rates in each cycle. Cycle 1 – all correct trials. Cycles 2 – 4, solid lines: correct trials, excluding cases with preceding error in this cycle. Cycle 2, dotted line: correct trial preceded by cycle 2 error. Cell selection and cross-validation (cycles 1 and 4 only) as for Figure 3.

In a supplementary analysis, we examined correct trials in cycle 2 that followed at least one cycle 2 error, a pattern suggesting failed learning on cycle 1 and continued exploration in cycle 2. For the two groups of explore cells, activity on these failed-learning trials resembled activity for cycle 1 (Figure 5, dotted green lines). Comparing failed-learning trials to regular cycle 2 trials, where the first response was correct (Figure 5, solid green lines), showed significant differences for both groups of cells (CH group, T = 3.66, P < .01; FB group, T = 3.19, P < .01). For exploit cells, results were less clear, with no significant difference between failed-learning and regular cycle 2 trials (CH group, T = 1.57; FB group, T = 0.79), and for FB cells, a significant difference between failed-learning and cycle 1 (T = 2.78, P < .01). These data show that, if learning was not complete after cycle 1, the frontal explore state was partially preserved into cycle 2.

### Expectancy and error

Slow, incremental learning is critically driven by reward prediction and prediction error. We wondered whether activity in our cell sample might be driven in part by prediction and error, factors that would strongly differentiate explore and exploit cycles. To test for such effects, we used activity in cycle 1, when the outcome of each selection was uncertain.

First, we asked how the activity of explore and exploit cells changed over the course of cycle 1, as more objects were sampled and eliminated, and the expectancy of reward progressively increased. For 1-target problems, we sorted correct (rewarded) cycle 1 trials according to whether the object selected was the first, second, third or fourth different object sampled in this cycle. For this analysis, it was impossible to use all display locations, because on the first trial, animals had very strong preferences for one location. To ensure that results were not biased by location preferences, we analyzed data just from trials in which the target was found in this same, favorite location, after sampling 0, 1, 2 or 3 other objects on previous trials (in any locations; note random repositioning of objects on each trial). Inconsistent with a progressive increase in reward expectancy, the results (SI Appendix, Figure S3) showed no significant effects of sampling order.

Second, we compared activity on correct (target selection) and incorrect (non-target selection) trials, again using cycle 1 data from 1-target problems. To eliminate effects of serial position in the cycle, we ignored data for revisits, and compared correct and incorrect trials unweighted for serial position (average of responses separately calculated for the first, second and third objects sampled in the cycle; incorrect impossible for object sampled fourth). Again the analysis was conducted just for objects in the animal’s preferred location. Though no difference between correct and incorrect selections could be expected at CH, these two sets of trials have very different reward prediction errors at FB. Even at FB, however, results suggested little difference between correct and incorrect trials (SI Appendix, Figure S4). Note that, in the cell sample as a whole, there was frequent discrimination of corrects and errors in the FB period. ANOVA with factors correct/error x object set x target location showed that, in the whole sample of 254 cells, there were 58 (22.8%) with a main effect of correct/error, 24 preferring correct and 34 preferring error. Of the 52 explore/exploit cells defined in our main analysis at FB, 16 (30.8%) also showed a significant difference between corrects and errors. Thus outcome information was encoded in prefrontal cells, but rather independent of explore/exploit selectivity.

Finally, we compared target-discovery trials in 1-target problems with first and second targets discovered in 2-target problems. Again, these cases have very different reward expectancies; for example, in a 1-target problem, the first object selected has only a 0.25 probability of being a target, while for a 2-target problem, this probability is 0.5. Again, however, results showed very similar responses for these 3 types of cycle 1 target trials (SI Appendix, Figure S5).

Contrary to incremental changes in reward prediction, these data show that explore/exploit selectivity was approximately binary, distinguishing simply an explore state, in which new information was sought, and an exploit state, in which known information was used.

### Parietal activity

Finally, parallel analyses were conducted on a population of 170 cells recorded in inferior parietal cortex (for details see (29)). In this case, cross-validated testing did not identify significant explore-exploit selectivity, though trends in the data (SI Appendix, Figure S6) weakly resembled those found in frontal cells.

## Discussion

While learning can be slow when being done from scratch, well developed internal task models may be generalized to a newly encountered situation. In this case, new stimuli can be quickly bound to their roles within the task model. To use such a task model, different computations are required during information seeking and information use, and for different task operations within each learning stage. To examine the transition from information seeking to information use, we employed a task showing rapid, often one-shot learning.

First, we observed different patterns of frontal activity for explore and exploit. Such differences were highly volatile within a trial: matching many reports of mixed selectivity in prefrontal cells (16, 17), including independent feature preferences in different phases of a trial (30, 31), we found that preferences for explore vs exploit were independent during CH and FB. Internal models for these two stages of the task would involve very different cognitive operations – for explore, hypothesis generation at CH and learning at FB, but for exploit, retrieval at CH and confirmation at FB. As recent modelling work shows that such orthogonal coding can support continuous learning of different task operations (32), Conjunctive coding for combinations of trial phase (CH, FB) and knowledge state (explore, exploit) may be required to construct and direct the multiple stages of the abstract task model, allowing newly-learned information to be bound to correct task operations and then used to control subsequent behavior.

Second, we examined changes in the pattern of frontal activity associated with one-trial learning. Matching behavior, the switch from explore to exploit patterns was close to binary, with little influence of continuous changes in reward prediction or reward prediction error. When learning was not complete after the first success trial, furthermore, the data showed substantial though not complete preservation of the frontal explore state. While reward prediction error is an integral part of many learning processes, additional processes may be critical in rapid, model-based role learning. Our results are reminiscent of recent work on deep reinforcement learning, where meta-learning can allow rapid hypothesis-driven experimentation to replace slow parameter tuning (33).

Evidently, a learned task model must govern the shift in frontal state that accompanies and may implement the switch from explore to exploit. An important open question is how this model is created. Modelling studies have examined how progressive learning in neural networks can shape connectivity to implement required cognitive operations (34, 35), and one possibility is that this progressive shaping occurs within prefrontal cortex. The present data, however, show only the final state of the network when a task is well known.

Many studies show closely similar neural properties in lateral prefrontal and inferior parietal cortex (36–39). In the current task, indeed, lateral prefrontal and inferior parietal neurons show similar coding of target identity and location (29). Compared to prefrontal cells, however, inferior parietal cells showed only weak, generally non-significant hints of explore-exploit preference. Within an abstract task model, prefrontal cortex may play the most crucial part in switching between learning and use of object-role bindings.

Previous findings from both human imaging (27, 28) and single cell physiology (23, 24) suggest a reduction in frontal activity with the transition from unknown to known task rules, or more broadly over the early trials of a new task (40). In contrast to this simple change, we observe cells with both increased and decreased activity with the switch from explore to exploit. Both explore and exploit preferences may be important to direct the different cognitive operations of constructing and using the task model.

In one-shot learning, newly acquired information is bound to its role within a previously-learned, abstract task model. Building on previous findings (23, 24), our data show a one-shot switch of firing rate in many prefrontal cells, matching one-shot behavioral learning. This switch of neural activity occurs independently at different stages of a trial, with their different required operations. The binary switch in frontal activity may enable one-shot switch between cognitive operations of information seeking and information use. More generally, such switches may allow the high-speed adaptability that characterizes much animal and human behavior.

## Materials and Methods

Details of subjects, equipment, recording methods and recording locations have been described elsewhere (29). In brief, data were recorded from 2 male rhesus monkeys, across a total of 56 daily sessions. Recordings used a semi-chronic microdrive system (Gray Matter Research), with one 32-channel array over lateral frontal cortex (Figure 1B), the primary focus of the current report, and another over parietal cortex. The experiments were performed in accordance with the Animals (Scientific Procedures) Act 1986 of the UK; all procedures were licensed by a Home Office Project License obtained after review by Oxford University’s Animal Care and Ethical Review committee, and were in compliance with the guidelines of the European Community for the care and use of laboratory animals (EUVD, European Union directive 86/609/EEC).

Arrangement of each session into problems and cycles is described in the main text (see Figure 1A). 1-target and 2-target problems were blocked in each session (mean of 69 1-target and 67 2-target problems per session). Additional cues reinforced the animal’s knowledge of when each cycle and each problem were completed (see (29)).

Details of events on each trial are illustrated in Figure 1A (bottom). Before the trial began, the screen showed a central white fixation point (FP) and a surrounding display of 4 black squares (each square 5.7 x 5.7 deg visual angle, centred 11.4 deg from fixation). To initiate trial events, the monkey was required to press and hold down the start key, and to acquire and hold central fixation (window 7.6 x 7.6 deg). At this point the FP turned red, and there was a wait period of 0.8 to 1.2 s, after which the black squares were replaced by a display (CH) of 4 choice objects. Following a further delay of 1.2 to 2.0 s, the FP changed to cyan (GO) to indicate that a response could be made. To indicate his choice, the animal released the start key and touched one of the objects (touch required within 1.8 s of GO). After the touch had been held for 0.35 to 0.45s, the selected object was replaced by either a green (correct target touch) or red (incorrect) square (feedback, FB), which remained for 0.3 s followed by an inter-trial display. If the touch was correct, a drop of soft food was delivered 0.05 to 0.15 s after FB offset. Once a trial had been initiated, it was aborted without reward if the monkey released the start key or broke fixation prior to GO. The trial was also aborted if, after an object had been touched, the touch was not maintained until FB.

To produce PSTHs (Figures 3, 5) we counted spikes in 100 ms windows, starting at a window centered around -200 ms from CH or FB and then shifting in +25 ms steps to a final window centered at +475 ms. Spike counts in each window were divided by an estimate of the cell’s mean activity, defined as mean activity across all conditions in the CH and FB ANOVAs used for cell selection. To create the PSTH for each cell, we de-weighted mean activity in each window for the influence of number of targets (1, 2) x object set (1, 2) x touched location (1-4).

For all analyses, we excluded problems in which animals failed to respond on 6 or more trials in a single cycle, suggesting poor task focus.

## Acknowledgments

This work was funded by the Medical Research Council (UK, programme SUAG/045.G101400), the Wellcome Trust (grant 101092/Z/13/Z), a JSPS Postdoctoral Fellowship for Research Abroad to KW, and a Gates Cambridge Trust scholarship to JA.

## Supplementary tables

**Table S1.** Cycle 1, separate data for each animal: Percentage of trials with re-selection of a previously sampled object, as a function of number already sampled.

| Already<br>sampled | Animal A |  | Animal B |  | Expected |
| --- | --- | --- | --- | --- | --- |
|  | *1-target*<br>*problems* | *2-target*<br>*problems* | *1-target*<br>*problems* | *2-target*<br>*problems* |  |
| **0** | 0% | 0% | 0% | 0% | 0% |
| **1** | 6.4% | 5.7% | 2.9% | 6.2% | 25% |
| **2** | 26.1% | 24.4% | 12.1% | 17% | 50% |
| **3** | 55.5% | 54.2% | 40.6% | 44% | 75% |

## Supplementary figures

**Figure S1.**
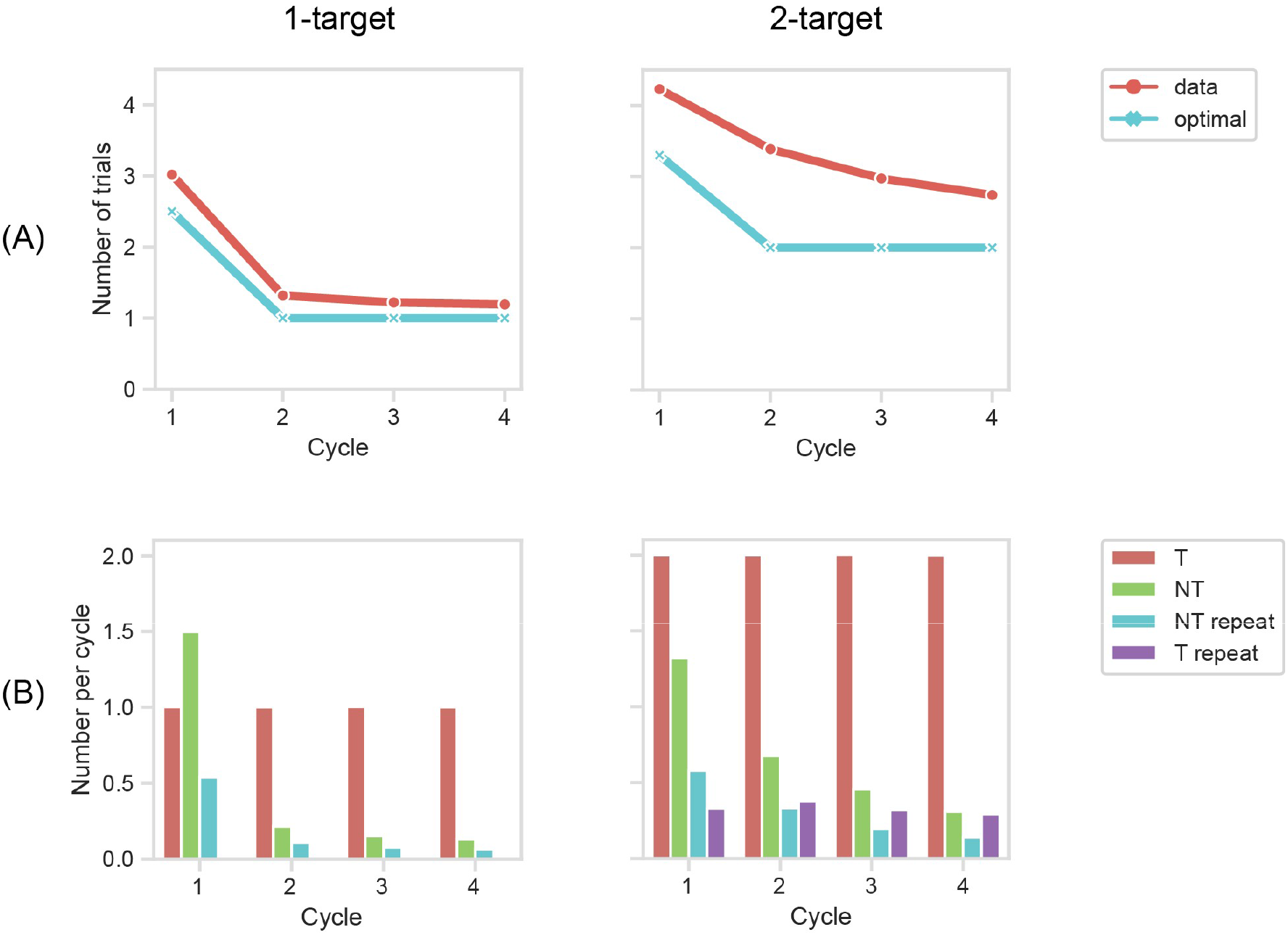
Behavioral data for animal A. (A) Mean number of trials per cycle. (B) Trials per cycle broken down into correct target selections (T), novel non-target selections (selection of a non-target not previously sampled in this cycle, NT), repeat non-target selections (selection of a non-target previously sampled in this cycle, NT repeat), and repeat target selections (T repeat; only possible for 2-target problems).

**Figure S2.**
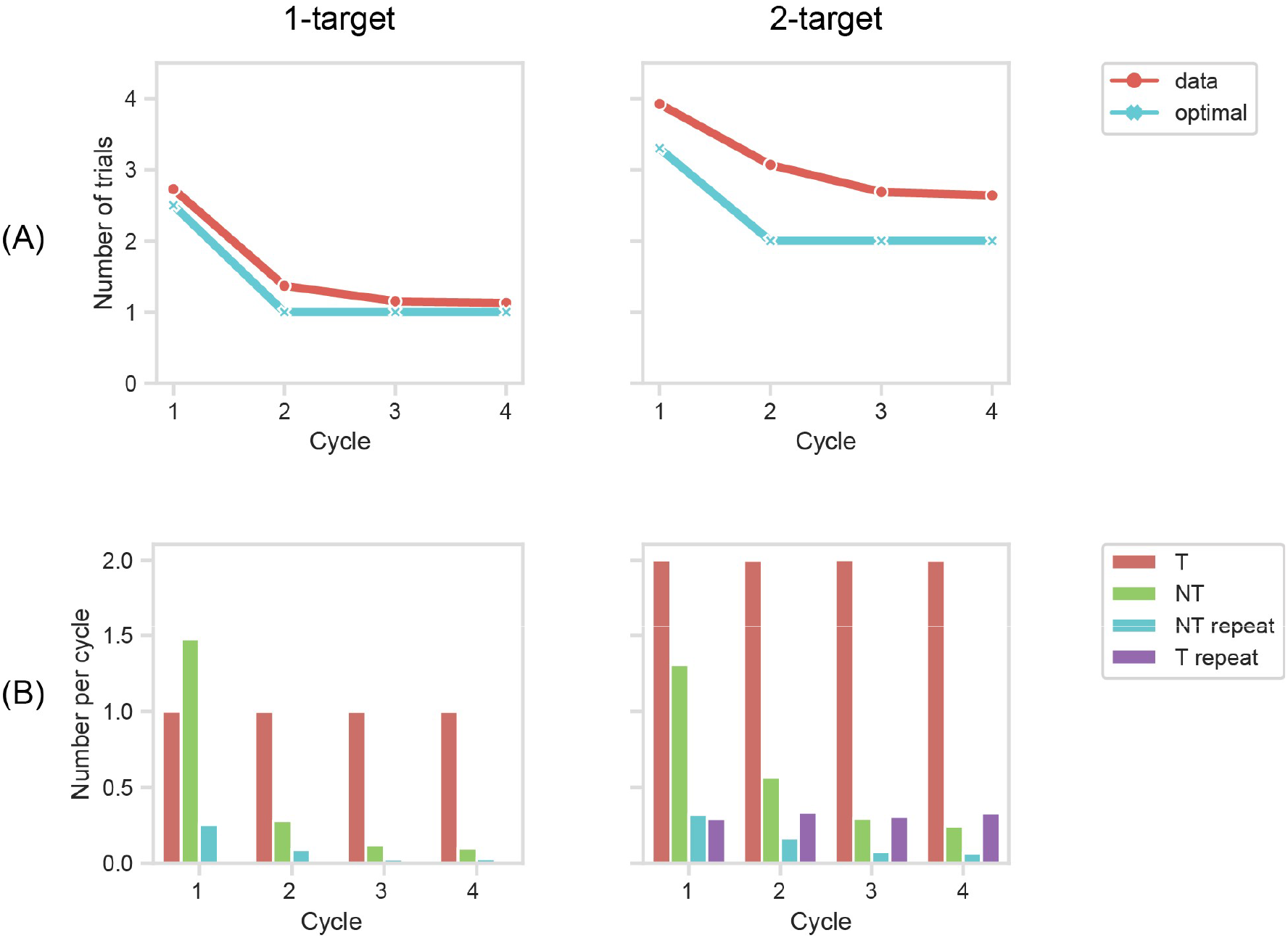
Behavioral data for animal B. (A) Mean number of trials per cycle. (B) Trials per cycle broken down into correct target selections (T), novel non-target selections (selection of a non-target not previously sampled in this cycle, NT), repeat non-target selections (selection of a non-target previously sampled in this cycle, NT repeat), and repeat target selections (T repeat; only possible for 2-target problems).

**Figure S3.**
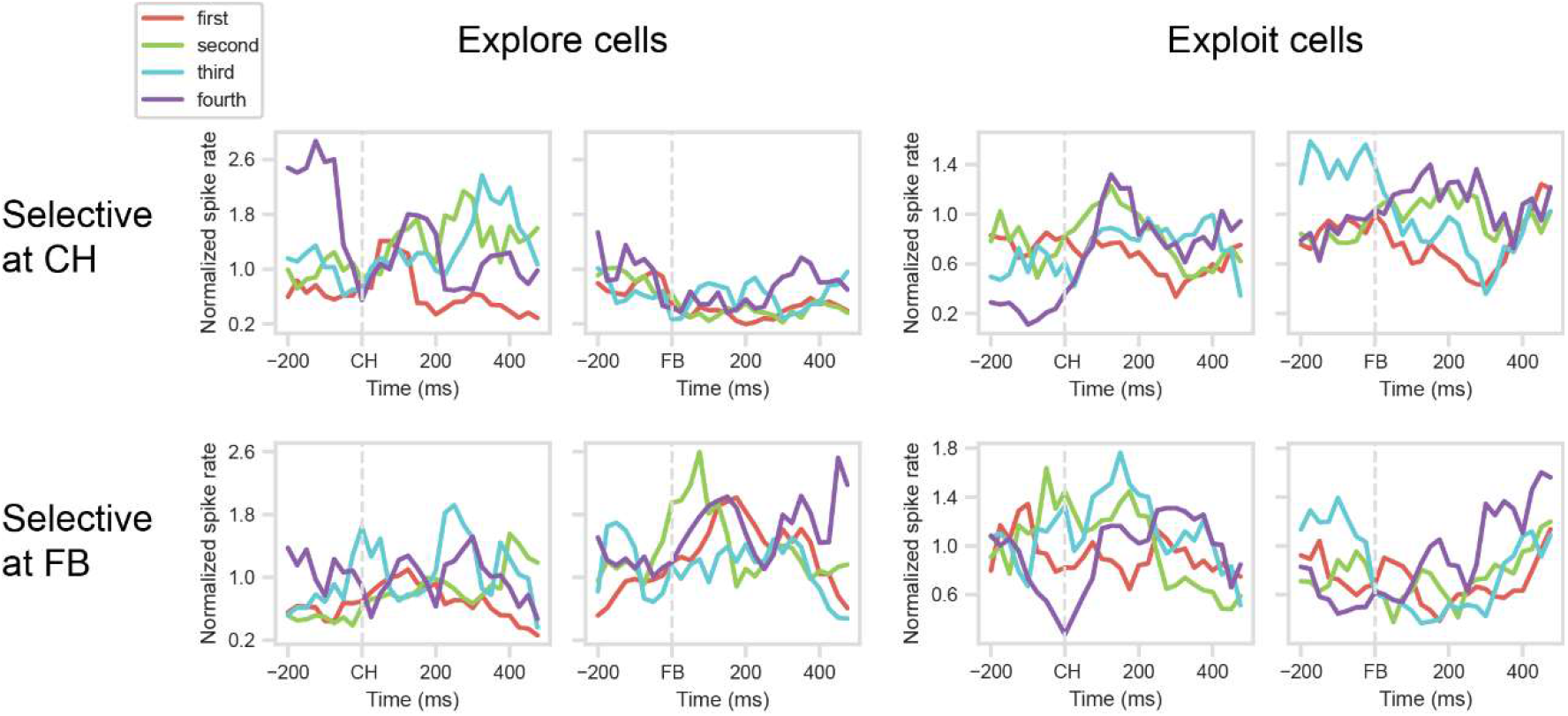
1-target-problems, cycle 1: Target first, second, third or fourth object sampled. Cell selection as Figure 3; data just for targets in animal’s preferred array location. For cells with an explore (cycle 1) or exploit (cycle 4) preference at CH (top row), ANOVAs on the 400 ms window following CH onset showed no significant differences between trials 1 – 4 (explore cells, F = 2.21; exploit cells, F = 0.88). For cells with an explore (cycle 1) or exploit (cycle 4) preference at FB (bottom row), similar results were obtained in the 400 ms window following FB onset (explore cells, F = 0.97; exploit cells, F = 1.70).

**Figure S4.**
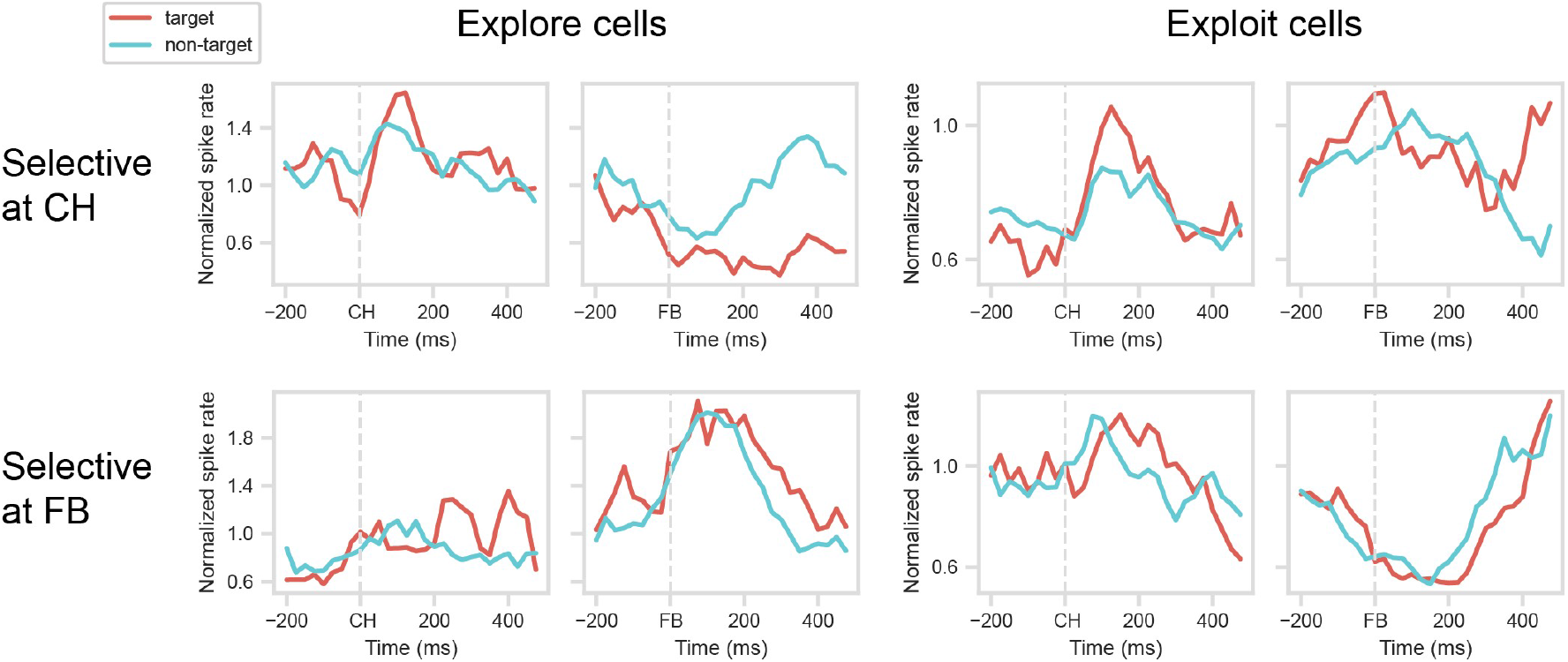
1-target-problems, cycle 1: Responses on correct (target selected) and incorrect (non-target selected) trials. Cell selection as Figure 3; data just for targets in animal’s preferred array location, averaged across first, second or third object sampled in cycle. For cells with an explore (cycle 1) or exploit (cycle 4) preference at CH (top row), T-tests on the 400 ms window following CH onset showed no significant differences between target and non-target trials (explore cells, T = 0.71; exploit cells, T = 0.84). For cells with an explore (cycle 1) or exploit (cycle 4) preference at FB (bottom row), similar results were obtained in the 400 ms window following FB onset (explore cells, T = 1.33; exploit cells, T = 0.96).

**Figure S5.**
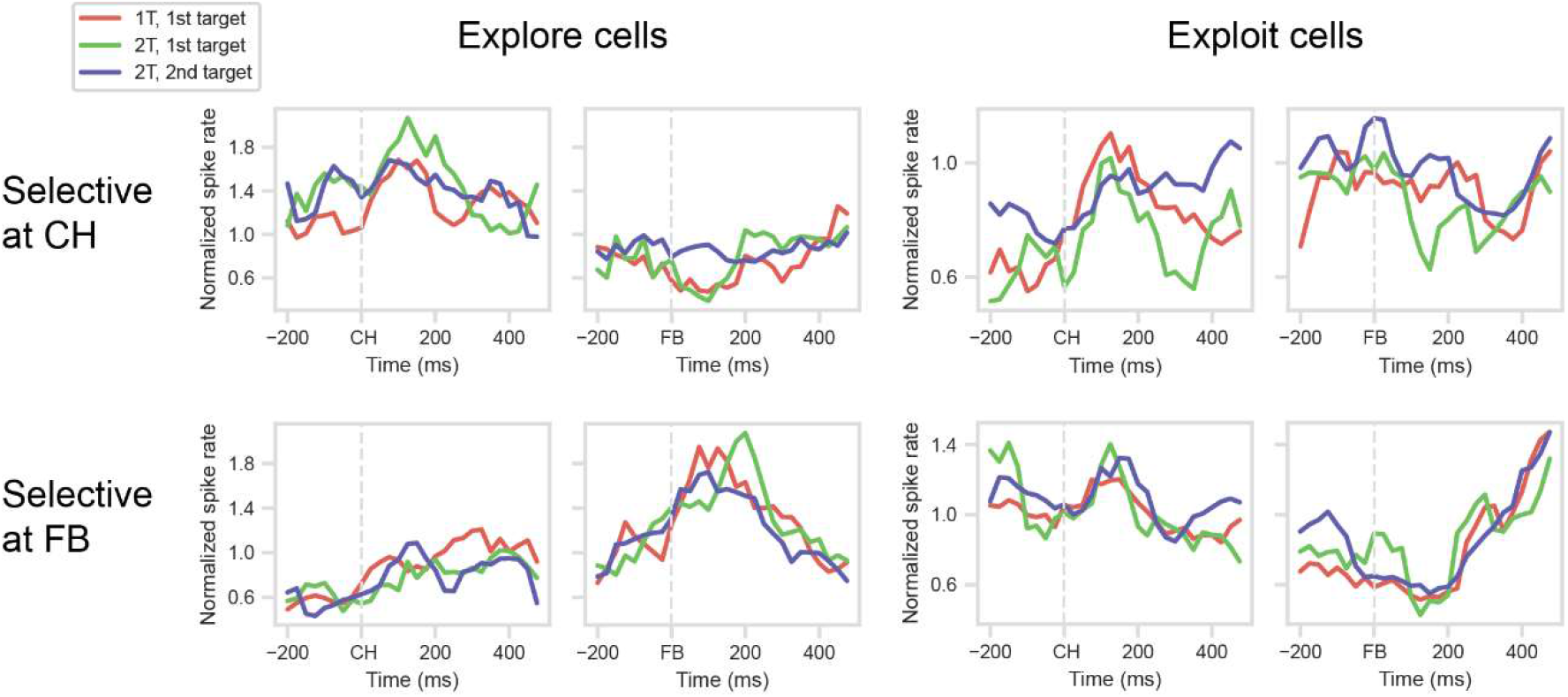
Cycle 1: comparison of 1-target problems with first and second targets discovered in 2-target-problems. Cell selection as Figure 3. For cells with an explore (cycle 1) or exploit (cycle 4) preference at CH (top row), ANOVAs on the 400 ms window following CH onset showed no significant differences between the three trial types (explore cells, F = 0.21; exploit cells, F = 2.78). For cells with an explore (cycle 1) or exploit (cycle 4) preference at FB (bottom row), similar results were obtained in the 400 ms window following FB onset (explore cells, F = 0.37; exploit cells, F = 0.25).

**Figure S6.**
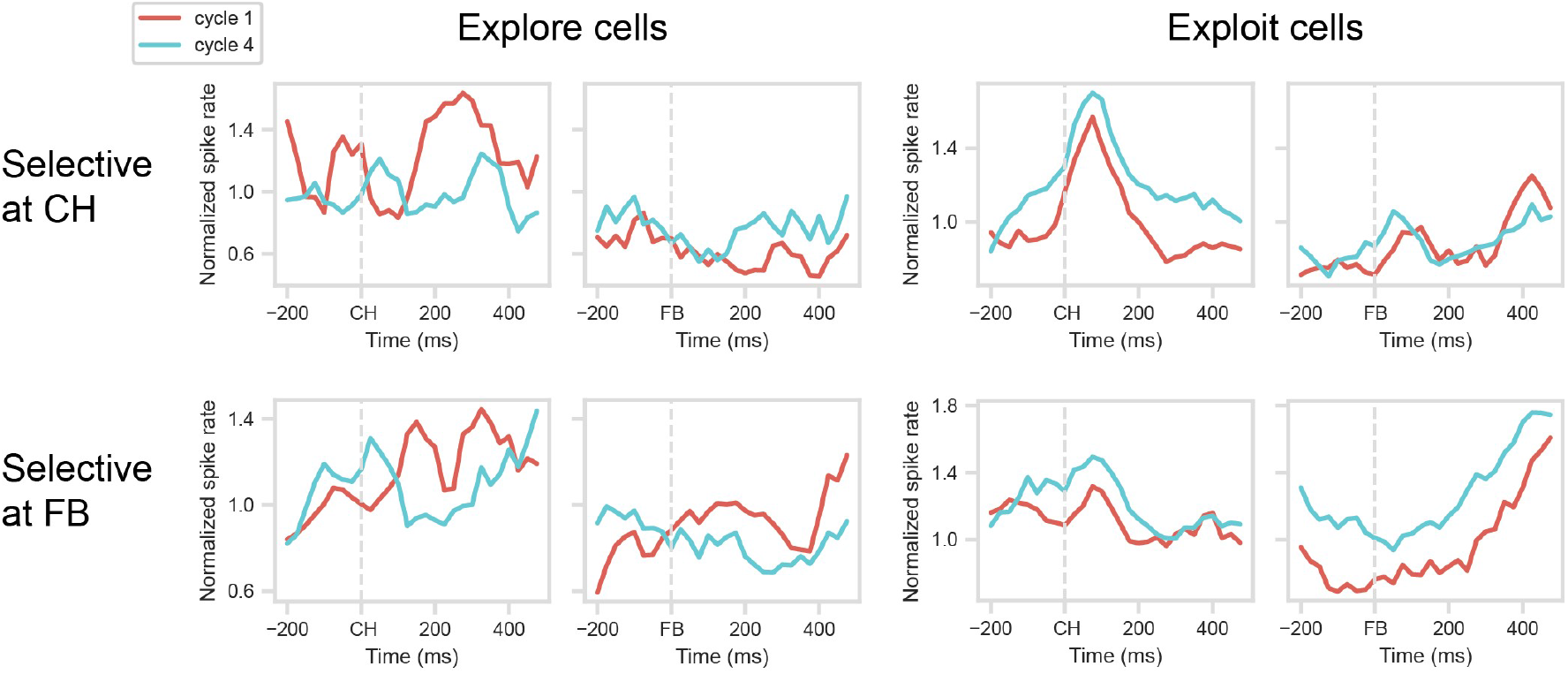
Data from inferior parietal cortex. Layout as Figure 3. For cells with an explore (cycle 1) or exploit (cycle 4) preference at CH (top row), T-tests on the 400 ms window following CH onset showed mixed results for differences between the cycles (explore cells, T = 0.80; exploit cells, T = 3.54, P < 0.001). For cells with an explore (cycle 1) or exploit (cycle 4) preference at FB (bottom row), similarly mixed results were obtained in the 400 ms window following FB onset (explore cells, T = 1.15; exploit cells, T = 3.65, P < 0.01).

## Notes

### Competing Interest Statement

The authors have declared no competing interest.

## References

1. R. A. Rescorla, A. R. Wagner, “A theory of pavlovian conditioning: Variations in the effectiveness of reinforcement and nonreinforcement” in Classical Conditioning II: Current Research and Theory, A. H. Black, W. F. Prokasy, Eds. (Appleton Century Crofts, 1972), pp. 64–99.

2. W. Schultz, P. Dayan, P. R. Montague, A neural substrate of prediction and reward. Science 275, 1593–1599 (1997).

3. J. Schmidhuber, Deep learning in neural networks: An overview. Neural Networks 61, 85–117 (2015).

4. Y. LeCun, Y. Bengio, G. Hinton, Deep learning. Nature 521, 436–444 (2015).

5. P. Smolensky, Tensor product variable binding and the representation of symbolic structures in connectionist systems. Artif. Intell. 46, 159–216 (1990).

6. H. F. Harlow, The formation of learning sets. Psychol. Rev. 56, 51–65 (1949).

7. T. E. J. Behrens, et al., What is a cognitive map? Organizing knowledge for flexible behavior. Neuron 100, 490–509 (2018).

8. J. X. Wang, et al., Prefrontal cortex as a meta-reinforcement learning system. Nat. Neurosci. 21, 860–868 (2018).

9. B. M. Lake, T. D. Ullman, J. B. Tenenbaum, S. J. Gershman, Building machines that learn and think like people. Behav. Brain Sci. 40, e253 (2017).

10. M. Rigotti, D. Ben Dayan Rubin, X.-J. Wang, S. Fusi, Internal representation of task rules by recurrent dynamics: The importance of the diversity of neural responses. Front. Comput. Neurosci. 4 (2010).

11. J. Duncan, M. Assem, S. Shashidhara, Integrated intelligence from distributed brain activity. Trends Cogn. Sci. 24, 838–852 (2020).

12. M. W. Cole, W. Schneider, The cognitive control network: Integrated cortical regions with dissociable functions. Neuroimage 37, 343–360 (2007).

13. M. Assem, M. F. Glasser, D. C. Van Essen, J. Duncan, A domain-general cognitive core defined in multimodally parcellated human cortex. Cereb. Cortex 30, 4361–4380 (2020).

14. E. K. Miller, J. D. Cohen, An integrative theory of prefrontal cortex function. Annu. Rev. Neurosci. 24, 167–202 (2001).

15. M. R. Warden, E. K. Miller, Task-dependent changes in short-term memory in the prefrontal cortex. J. Neurosci. 30, 15801–15810 (2010).

16. H. Mushiake, N. Saito, K. Sakamoto, Y. Itoyama, J. Tanji, Activity in the lateral prefrontal cortex reflects multiple steps of future events in action plans. Neuron 50, 631–641 (2006).

17. M. Rigotti, et al., The importance of mixed selectivity in complex cognitive tasks. Nature 497, 585–590 (2013).

18. J. M. Fuster, R. H. Bauer, J. P. Jervey, Cellular discharge in the dorsolateral prefrontal cortex of the monkey in cognitive tasks. Exp. Neurol. 77, 679–694 (1982).

19. E. K. Miller, C. A. Erickson, R. Desimone, Neural mechanisms of visual working memory in prefrontal cortex of the macaque. J. Neurosci. 16, 5154–5167 (1996).

20. F. A. Mansouri, Prefrontal Cell Activities Related to Monkeys’ Success and Failure in Adapting to Rule Changes in a Wisconsin Card Sorting Test Analog. J. Neurosci. 26, 2745–2756 (2006).

21. D. Durstewitz, N. M. Vittoz, S. B. Floresco, J. K. Seamans, Abrupt transitions between prefrontal neural ensemble states accompany behavioral transitions during rule learning. Neuron 66, 438–448 (2010).

22. R. Bartolo, B. B. Averbeck, Prefrontal cortex predicts state switches during reversal learning. Neuron 106, 1044–1054 (2020).

23. E. Procyk, P. S. Goldman-Rakic, Modulation of dorsolateral prefrontal delay activity during self-organized behavior. J. Neurosci. 26, 11313–11323 (2006).

24. R. Quilodran, M. Rothé, E. Procyk, Behavioral shifts and action valuation in the anterior cingulate cortex. Neuron 57, 314–325 (2008).

25. M. Rothe, R. Quilodran, J. Sallet, E. Procyk, Coordination of high gamma activity in anterior cingulate and lateral prefrontal cortical areas during adaptation. J. Neurosci. 31, 11110–11117 (2011).

26. M. Khamassi, R. Quilodran, P. Enel, P. F. Dominey, E. Procyk, Behavioral regulation and the modulation of information coding in the lateral prefrontal and cingulate cortex. Cereb. Cortex 25, 3197–3218 (2015).

27. A. Hampshire, A. M. Owen, Fractionating attentional control using event-related fMRI. Cereb. Cortex 16, 1679–1689 (2005).

28. S. Konishi, et al., Transient activation of inferior prefrontal cortex during cognitive set shifting. Nat. Neurosci. 1, 80–84 (1998).

29. M. Kadohisa, K. Watanabe, M. Kusunoki, M. J. Buckley, J. Duncan, Focused representation of successive task episodes in frontal and parietal cortex. Cereb. Cortex 30, 1779–1796 (2020).

30. N. Sigala, M. Kusunoki, I. Nimmo-Smith, D. Gaffan, J. Duncan, Hierarchical coding for sequential task events in the monkey prefrontal cortex. Proc. Natl. Acad. Sci. 105, 11969–11974 (2008).

31. M. G. Stokes, et al., Dynamic coding for cognitive control in prefrontal cortex. Neuron 78, 364–375 (2013).

32. L. Duncker, L. N. Driscoll, K. V Shenoy, M. Sahani, D. Sussillo, Organizing recurrent network dynamics by task-computation to enable continual learning in Advances in Neural Information Processing Systems 33 Pre-Proceedings (NeurIPS 2020), (Conference on Neural Information Processing Systems, 2020).

33. M. Botvinick, J. X. Wang, W. Dabney, K. J. Miller, Z. Kurth-Nelson, Deep reinforcement learning and its neuroscientific implications. Neuron 107, 603–616 (2020).

34. O. Barak, D. Sussillo, R. Romo, M. Tsodyks, L. F. Abbott, From fixed points to chaos: Three models of delayed discrimination. Prog. Neurobiol. 103, 214–222 (2013).

35. W. Chaisangmongkon, S. K. Swaminathan, D. J. Freedman, X.-J. Wang, Computing by robust transience: How the fronto-parietal network performs sequential, category-based decisions. Neuron 93, 1504-1517.e4 (2017).

36. M. V. Chafee, P. S. Goldman-Rakic, Matching patterns of activity in primate prefrontal area 8a and parietal area 7ip neurons during a spatial working memory task. J. Neurophysiol. 79, 2919–2940 (1998).

37. S. J. Goodwin, R. K. Blackman, S. Sakellaridi, M. V. Chafee, Executive control over cognition: Stronger and earlier rule-based modulation of spatial category signals in prefrontal cortex relative to parietal cortex. J. Neurosci. 32, 3499–3515 (2012).

38. E. M. Meyers, A. Liang, F. Katsuki, C. Constantinidis, Differential processing of isolated object and multi-item pop-out displays in LIP and PFC. Cereb. Cortex 28, 3816–3828 (2018).

39. S. L. Brincat, M. Siegel, C. von Nicolai, E. K. Miller, Gradual progression from sensory to task-related processing in cerebral cortex. Proc. Natl. Acad. Sci. 115, E7202–E7211 (2018).

40. H. Ruge, T. A. J. Schäfer, K. Zwosta, H. Mohr, U. Wolfensteller, Neural representation of newly instructed rule identities during early implementation trials. Elife 8 (2019).

